# A discrete protein subset drives structure prediction discordance in orphan proteins

**DOI:** 10.64898/2026.08.24.746756

**Authors:** Lars A. Eicholt, Lasse Middendorf

## Abstract

Structure and disorder predictors are increasingly used as decision-grade tools in protein engineering and in the analysis of newly emerged proteins, yet how the current state-of-the-art behaves on sequences outside the well-charted evolutionary space remains poorly characterised. We previously reported that AlphaFold2 confidence and the disorder predictor flDPnn produced discordant predictions for naturally evolved *de novo Drosophila* proteins and for shuffled sequences. Here, we revisit the comparison with AlphaFold3 and the best-performing disorder predictor PUNCH2 on the same sequence sets together with conserved *Drosophila* proteins and intrinsically disordered proteins. The discordance persists: pLDDT correlates positively with PUNCH2 disorder in random and *de novo* proteins and negatively with *β*-strand fraction, opposite to the conserved and disordered baselines. A class-specific, score-defined driver subset jointly captures the unusual high-pLDDT, high-disorder, low-strand combination and contains 24.5% of *de novo*, 29.4% of random, 5.1% of conserved, and 1.3% of disordered proteins. Removing this subset normalises the correlations. A held-out classifier trained on architectural and compositional features that were *not* used in the driver definition recovers the subset, with helix and coil fraction, sequence length, entropy and hydropathy as the strongest predictors. The discordance is therefore not a sequence-class artefact but a localised, compositionally identifiable phenotype that current predictors handle in a non-canonical way - a concrete failure mode that protein designers and other working on sequences remote in sequence space should be aware of when relying on predictor outputs.

## 1 Introduction

The combination of experimentally solved protein structures ^1^, co-evolution based inference of structural proximity between amino acid residues ^2–9^, and modern machine-learning architectures ^10–15^ combined with hardware advances ^16,17^, has driven a renaissance in protein structure prediction and design ^18^. AlphaFold2^19^ reached for the first time experimental accuracy on most globular domains in 2021 assessed during CASP 14 (Critical Assessment of protein Structure Prediction; a bi-annual competition of protein structure prediction tools) ^20^, and was followed by AlphaFold3, which extends modelling to a broad range of biomolecular complexes including nucleic acids, ligands, and post-translational modifications ^21^. In parallel, intrinsic disorder prediction has matured around the Critical Assessment of Intrinsic Disorder (CAID) benchmark ^22^, which has identified successively accurate predictors such as flDPnn ^23^ and, in the most recent third iteration of CAID ^24^, identified PUNCH2 / Punch2-light ^25^, built on protein language model embeddings ^26^, as the best disorder predictor. Nevertheless, these tools have a strong dependency on evolutionary signals. AlphaFold-style models exploit co-evolving residues identified in MSAs and rely indirectly on the experimentally solved structures used during training ^19,21^. Sequences for which deep MSAs cannot be built or experimental structures are limited ^27^ - including computationally designed *de novo* proteins, randomly shuffled sequences, and naturally emerged *de novo* proteins originating from previously non-coding DNA ^28^ - are notoriously difficult targets for computational predictions. Protein language models (pLMs) have been proposed to perform better on such orphan sequences - sequences lacking similarity to other sequences ^29^ but their performance is inherently bound to how close the orphan sequences tested are to the sequences the pLM was trained on ^30^. Hence, sequences lacking similarity to any other, probe machine-learning based structure and disorder predictors at the limits of their training distribution ^31–35^. Reliable structure prediction in these regimes is increasingly load-bearing: pLM based and AlphaFold-style scoring functions are central to *de novo* design pipelines, where the encoded sequence is checked back against the intended backbone using exactly these structure predictors ^36–39^. Disorder predictors have similarly become a standard analytical tool in the field of *de novo* emerged proteins ^31^, and the choice of predictors and thresholds has been actively debated ^31,40^. Additionally, AlphaFold predictions have now become a frequently used tool in studies on *de novo* emerged proteins ^41–51^. Yet the performance of such predictors on sequences outside well-charted evolutionary space remains poorly characterised ^52^.

In our previous study ^32^ we used AlphaFold2^53^ and flDPnn ^23^, the then state-of-the-art tools, to characterise naturally emerged *de novo* proteins from twelve *Drosophila* species ^54^, randomised (amino-acid shuffled) controls matched in length and amino-acid composition, a conserved *Drosophila* reference set ^32^, and a set of experimentially confirmed intrinsically disordered proteins (IDPs) ^55^. We found that for *de novo* and random sequences, mean prediction confidence (predicted local distance difference test; pLDDT ^19,56^) correlated negatively with *β*-sheet content and positively with predicted disorder, opposite to the canonical relationship observed for well-folded conserved proteins ^32^. We were also able reproduce those discordances using pLM-based structure predictor ESMfold ^57^ and Iupred3^58^. Whether this discordance was a property of those specific predictors or a more general signal of how current models handle sequences outside their training distribution has remained open.

Here we revisit the question with the current best-performing structure and disorder predictors, AlphaFold3^21^ and PUNCH2^25^, on the same four sequence classes: 2,506 *de novo Drosophila* proteins, 2,507 length- and composition-matched random sequences, 2,224 conserved *Drosophila* proteins, and 2,066 experimentally validated intrinsically disordered proteins from DisProt ^55^. We omit protein language model based structure predictors such as ESMFold because (i) we and others previously reported the same discordant behaviour ^31,32,48^ and (ii) these models did not match AlphaFold-style accuracy in CASP15 despite their relative independence from MSAs ^59^. We chose *Drosophila de novo* proteins because their non-coding ancestry has been explicitly verified ^54^ and because they span a useful length distribution; alternative datasets such as *Saccharomyces cerevisiae de novo* candidates are more strongly biased toward shorter sequences ^60,61^ - with short sequence length being a known caveat for structure and disorder predictors ^62–65^

We first confirm that the discordance persists for AlphaFold3 and PUNCH2. We then use a class-specific score combining pLDDT, disorder, and strand content to define a subset of proteins that jointly drive the unusual correlations (high pLDDT for high disorder; low pLDDT for *β*-sheets). Removing this subset restores the canonical correlation pattern (low pLDDT for high disorder; high pLDDT for *β*-sheets). We test, and rule out, transmembrane helices - which are common in *de novo* emerged proteins ^33,66,67^ and can be transient between structural and disorder states ^68^ - as the dominant mechanistic driver, and we show that an independent classifier trained on non-defining architectural and compositional features recovers the same subset. To allow others to test their predictions easily for discordance, we provide a driver assessment command-line tool (see Data and software availability). We close by discussing the implications for protein design and *de novo* gene emergence research, where structure and disorder predictors are increasingly used as decision-grade tool on sequences far out from the well-charted evolved sequence space ^69^.

## 2 Materials and Methods

### 2.1 Sequence datasets

The sequence datasets were generated in Middendorf & Eicholt (2024). Briefly, sequences whose mechanism of emergence was annotated as *de novo* or *de novo*-intronic in Heames *et al*. (2020) were retained, yielding 2,506 non-redundant *de novo Drosophila* protein sequences (de_novo). Length- and composition-matched random sequences (2,507 unique sequences; random) were generated using the build_oligos.py script of Heames *et al*. (2023), which samples amino acids according to the empirical *de novo* amino-acid frequency under the constraint that the first residue is methionine. Conserved control proteins were sampled from the combined proteome of eleven *Drosophila* species (*D. simulans* (7220), *D. melanogaster* (7227), *D. persimilis* (7234), *D. sechellia* (7238), *D. virilis* (7244), *D. mojavensis* (7245), *D. erecta* (7230), *D. grimshawi* (7222), *D. pseudoobscura* (7237), *D. willistoni* (7240), and *D. yakuba* (7260)) to match the per-protein length distribution of the *de novo* set; after removal of duplicates and *de novo* entries, this gave 2,224 conserved sequences (conserved). Experimentally validated intrinsically disordered proteins were taken from DisProt release 12/2022^55^ (2,066 sequences; disprot). All datasets are deposited on Zenodo (Data Availability). DeNoFo annotation file ^51,71^ is deposited in Zenodo.

### 2.2 Structural and disorder predictions

Structure predictions were locally performed with AlphaFold3^21^; for each sequence, the model with the highest mean pLDDT (predicted local distance difference test) was retained. Per-residue secondary structure was annotated with the pydssp re-implementation of DSSP ^72,73^; the helix (H), strand (E) and coil (C) fractions reported here are calculated as the number of residues in the corresponding DSSP state divided by sequence length. Disorder was predicted with Punch2-light ^25^ using ProtTrans ^26^ and one-hot embeddings; residues with PUNCH2 score ≥ 0.5 were defines as disordered. Transmembrane (TM) and signal-peptide propensity were predicted with Phobius ^74^. All per-protein and per-residue predictions are deposited on Zenodo.

### 2.3 Driver definition

To identify proteins driving the positive correlation between mean predicted disorder and mean pLDDT, together with the negative correlation between strand fraction and mean pLDDT, we defined a protein-level *driver score*:

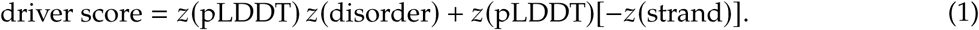

where *z*(·) denotes within-class standardisation. Proteins with driver score ≥ 1.0 were classified as drivers. This definition was used as the primary operational definition of driver proteins throughout the analyses.

### 2.4 Driver-subset analyses

Class-specific Spearman correlations between mean pLDDT, mean disorder, helix fraction and strand fraction were recomputed on (i) all proteins, (ii) drivers only, and (iii) non-drivers. Additional analyses re-evaluated the correlations after excluding TM residues, helical residues, or TM-helical residues. Continuous protein- or segment-level properties were compared between drivers and non-drivers using Mann-Whitney *U* tests ^75^ with Benjamini-Hochberg false-discovery-rate correction ^76^; Cliff’s *δ* was reported as the effect size ^77,78^. Within-protein paired comparisons of state-specific mean pLDDT (e.g. helix vs. coil within the same protein) were assessed by Wilcoxon signed-rank tests ^79^. Binary enrichment was tested with Fisher’s exact test ^80^.

To test whether score-defined drivers represent distinct subtypes, drivers were post-hoc clustered on standardised non-defining structural and compositional features (helix and TM fractions, signal-peptide fraction, low-complexity and repeat fractions, hydropathy, charge, sequence length, regional helix/TM/disorder fractions, selected amino-acid frequencies); subtype labels were assigned from centroid properties. To capture local rather than protein-wide signatures, residue windows of fixed length (15 residues) were analysed. Window-level local-driver states were defined within each class using the same score logic as the protein-level definition combined with above-median window pLDDT and disorder, and enrichment of helix-rich, strand-rich, TM-rich, disorder-rich, and terminal versus central windows was tested against the remaining windows. To investigate local sequence grammar, class-specific 3-mer and 5-mer enrichment analyses were performed within helical windows, strand-rich windows, and windows with mean disorder score > 0.5. For each context, driver and non-driver *k*-mer counts were compared using Fisher’s exact tests with Benjamini-Hochberg correction, and effect sizes were summarised as log_2_ odds ratios. Helix-focused analyses were additionally performed at the segment level: contiguous AlphaFold3-predicted helical segments were extracted per protein and compared between drivers and non-drivers for length, mean pLDDT, mean disorder, mean hydropathy, TM overlap, low-complexity overlap, repeat overlap, hydrophobic moment, and amino-acid group composition.

### 2.5 Driver classifier

The classifier was *not* used to define drivers. Instead, it asked whether the score-defined driver subset is also recoverable from non-defining features alone, i.e. whether the subset occupies a separable region of architecture and composition space. For each class, we trained class-specific supervised classifiers on protein-level non-defining features only: helix and coil fractions; transmembrane and signal-peptide summaries; low-complexity and homorepeat fractions; Shannon entropy; hydropathy; charge; sequence length; regional helix/coil/TM/disorder fractions in N-, central-, and C-terminal thirds; per-amino-acid frequencies. Variables directly used in, or closely correlated with, the driver definition (mean pLDDT, mean disorder, strand fraction and derivatives) were excluded.

Initial benchmarking was performed within each class using nested stratified cross-validation. The outer loop split the data into three stratified folds for evaluation; within each outer training split, an inner three-fold stratified cross-validation procedure was used for model selection. We compared a dummy classifier (class prevalence only), *L*_2_-regularised logistic regression, histogram gradient boosting (HGB) ^81^, and a multilayer perceptron ^82^. Logistic-regression features were standardised within each training fold using StandardScaler (scikit-learn 1.7.0); HGB models were trained on the same non-defining feature sets without feature scaling. Held-out performance was summarised using average precision, ROC AUC, and Brier score. To test robustness against feature leakage and circularity, classifier performance was re-evaluated under progressively stricter non-defining feature sets (a strict full non-defining set, an ultra-strict set with additional exclusions, an architecture-only set, and a composition-only set), and label-permutation controls were run to verify that predictive performance exceeded chance.

HGB was the most stable family across the strict non-defining sets and was retained as the final model with learning rate 0.05, maximum depth 3, 150 boosting iterations, and minimum leaf size 20. Final performance was estimated by repeated stratified 3-fold cross-validation with 20 repeats; held-out out-of-fold predictions were averaged across repeats to yield a per-protein mean repeated out-of-fold driver probability. Average precision, ROC AUC, precision, recall and F1 at probability threshold 0.5 are reported. Mean ROC and precision-recall curves were computed by averaging fold-wise curves across repeats. Held-out permutation importance was computed by permuting each feature in the test fold, measuring the drop in average precision, and averaging across folds and repeats. Dimensionality-reduction analyses (PCA and UMAP) were repeated on standardised non-defining features, with points coloured by mean repeated out-of-fold driver probability. Score-defined and classifier-recognised drivers were compared in terms of overlap counts within each class.

### User-facing driver assessment tool

To facilitate external use of the published score-defined driver rule, we provide a separate command-line tool, 31_assess_driver_candidates.py, for assessment of one or more user-supplied protein structures. The script accepts individual structure files or AlphaFold3 output directories, extracts mean pLDDT-like confidence values from the structure, matches PUNCH2-style disorder predictions, and evaluates each input using the same driver-score definition as in the main analysis: *z*(mean pLDDT) × *z*(mean disorder) + *z*(mean pLDDT) × [−*z*(strand fraction)], with within-class standardization against the reference protein sets and the same threshold of ≥ 1.0. Secondary-structure features can be provided either via a supplied DSSP-derived summary table or by on-the-fly DSSP assignment, which uses pydssp when available and otherwise falls back to mkdssp. The utility reports class-specific driver scores, whether the published threshold is reached in any reference class context, and whether a result is indeterminate because disorder or strand-fraction estimates are unavailable. Sequence-only input is not supported because the workflow depends on structure-derived pLDDT-like values and secondary-structure assignment.

### 2.6 Software

Analyses were run in Python 3.13.1 with Pandas 2.2.3^83^, NumPy 2.2.2^84^, SciPy 1.15.1^85^, statsmodels 0.14.4^86^, scikit-posthocs 0.11.4^87^, scikit-learn 1.7.0^88^, Biopython 1.85^89^, Matplotlib 3.10.0^90^, Seaborn 0.13.2^91^, gemmi 0.7.3^92^, pyarrow 23.0.1^93^, pynndescent 0.5.13^94^, umap-learn 0.5.9^95^, and numba 0.61.2^96^.

## 3 Results

### 3.1 Class-level distributions of pLDDT, disorder, and secondary structure

We first asked whether the four sequence classes display differences in AlphaFold3 confidence, PUNCH2 disorder, and DSSP secondary-structure composition (Figure 1; Supplementary Figures S1-S8). Mean pLDDT was highest for DisProt (mean 73.6, median 75.4) and conserved (mean 70.0, median 70.1) proteins, intermediate for *de novo* (mean 52.9, median 52.6) and lowest for random sequences (mean 45.7, median 44.9); all pairwise differences except DisProt vs. random were significant (Mann-Whitney *U* test, all *q* ≤ 1.3 × 10^−20^, Cliff’s *δ* for conserved vs. random = 0.84, Kruskal-Wallis *η*^2^ = 0.49). PUNCH2 disorder was highest in *de novo* (mean 0.66), intermediate in random (0.50) and conserved (0.44), and *lowest* in DisProt (0.38); the apparent paradox that DisProt has lower mean PUNCH2 disorder than *de novo* reflects the size of DisProt entries (median 462 residues vs. 103-126 for the other classes), which dilutes annotated disordered regions. Helix fraction was highest in conserved (mean 0.43) and lowest in random (mean 0.28, *δ* = −0.33 vs. conserved, *q* = 4 × 10^−86^); strand fraction was highest in DisProt (mean 0.13, with substantial *β*-rich subsets) and lowest in *de novo* (mean 0.06). Together, these distributions confirm that the four classes are distinct in their distributions of structural elements and not only in their central tendencies (Supplementary Figures S1-S8).

**Figure 1:**
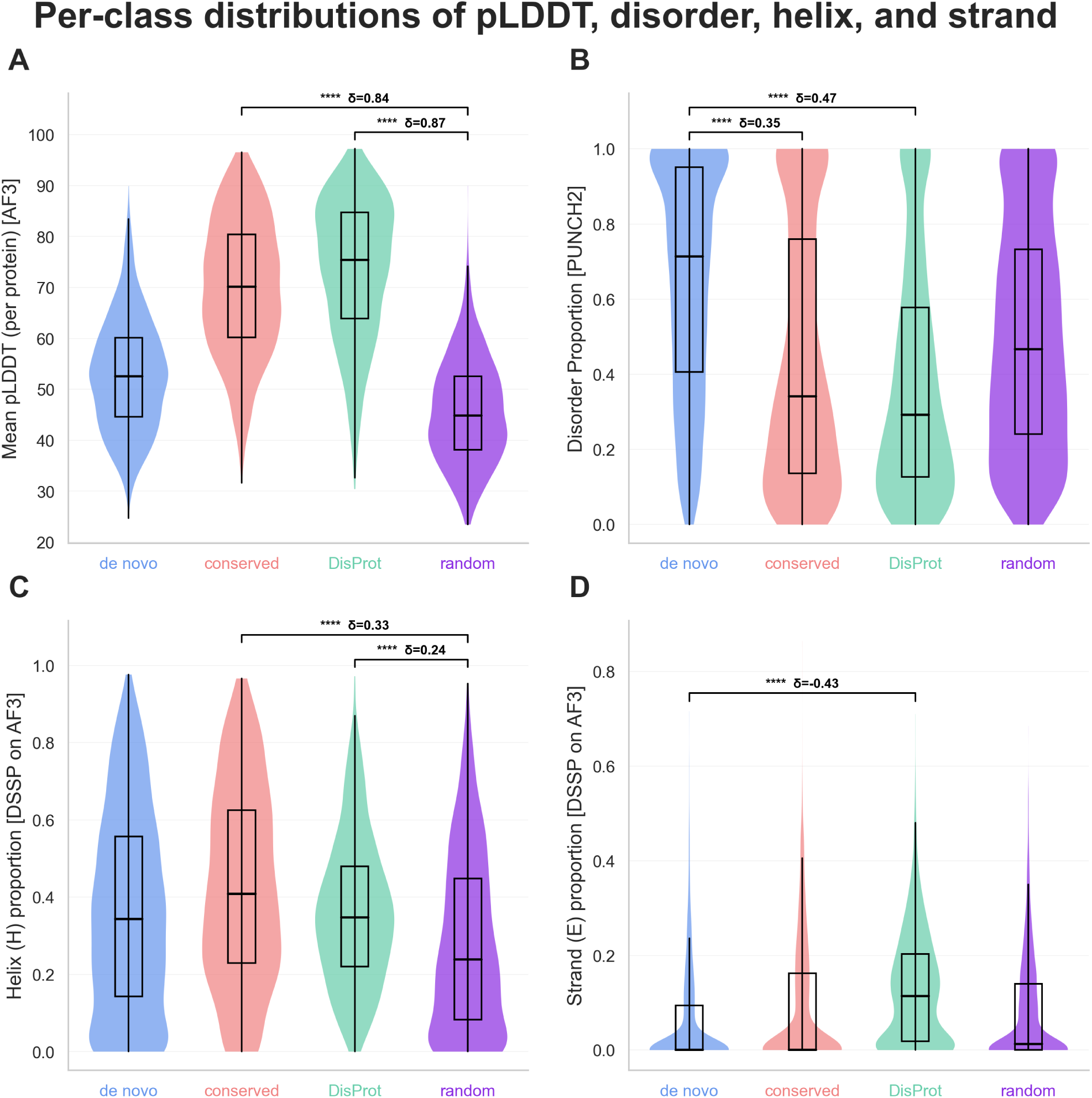
Class-level distributions of AlphaFold3 confidence, PUNCH2 disorder and DSSP secondary structure. **(A)** Mean per-protein AlphaFold3 pLDDT for *de novo*, conserved, DisProt and random proteins. **(B)** Mean per-protein PUNCH2 disorder proportion. **(C)** Fraction of residues assigned to helix state H by DSSP. **(D)** Fraction of residues assigned to strand state E by DSSP. Violin plots show the full per-protein distributions, with embedded box plots indicating median and interquartile range. Brackets mark selected pairwise comparisons; asterisks indicate significance level and *δ* denotes Cliff’s effect size.

### 3.2 pLDDT, disorder, and secondary-structure relationships are class-specific and discordant for random and *de novo* sequences

The expected canonical pattern - high pLDDT couples to low disorder, more *β*-sheets, and *α*-helices - holds for conserved and DisProt proteins. Mean pLDDT was negatively correlated with PUNCH2 disorder in conserved (*ρ* = −0.60, *q* = 6 × 10^−220^) and very strongly so in DisProt (*ρ* = −0.85, *q* < 10^−300^), and positively correlated with helix and strand fractions (*ρ*_helix_ = +0.24 and +0.34; *ρ*_strand_ = +0.27 and +0.40, respectively; Figure 2). For random and *de novo* sequences, the pLDDT-disorder relationship was inverted (random: *ρ* = +0.33, *q* = 5 × 10^−65^; *de novo*: *ρ* = +0.12, *q* = 1 × 10^−9^), and pLDDT-strand was *negative* (random: *ρ* = −0.39, *q* = 2 × 10^−92^; *de novo*: *ρ* = −0.43, *q* = 2 × 10^−111^). Helix-fraction-pLDDT remained positive across all four classes (*ρ* ≥ +0.24). Region-stratified, calibration, and partial-correlation summaries (Supplementary Figures S9-S17, S22, S29, S30) show that the inverted signs are robust to N-/centre-/C-terminal stratification and to partialling out hydropathy, complexity, and amino-acid composition. We therefore confirm with the current best-of-class predictors AlphaFold3 and PUNCH2 the discordance previously reported for AlphaFold2 and flDPnn ^32^.

**Figure 2:**
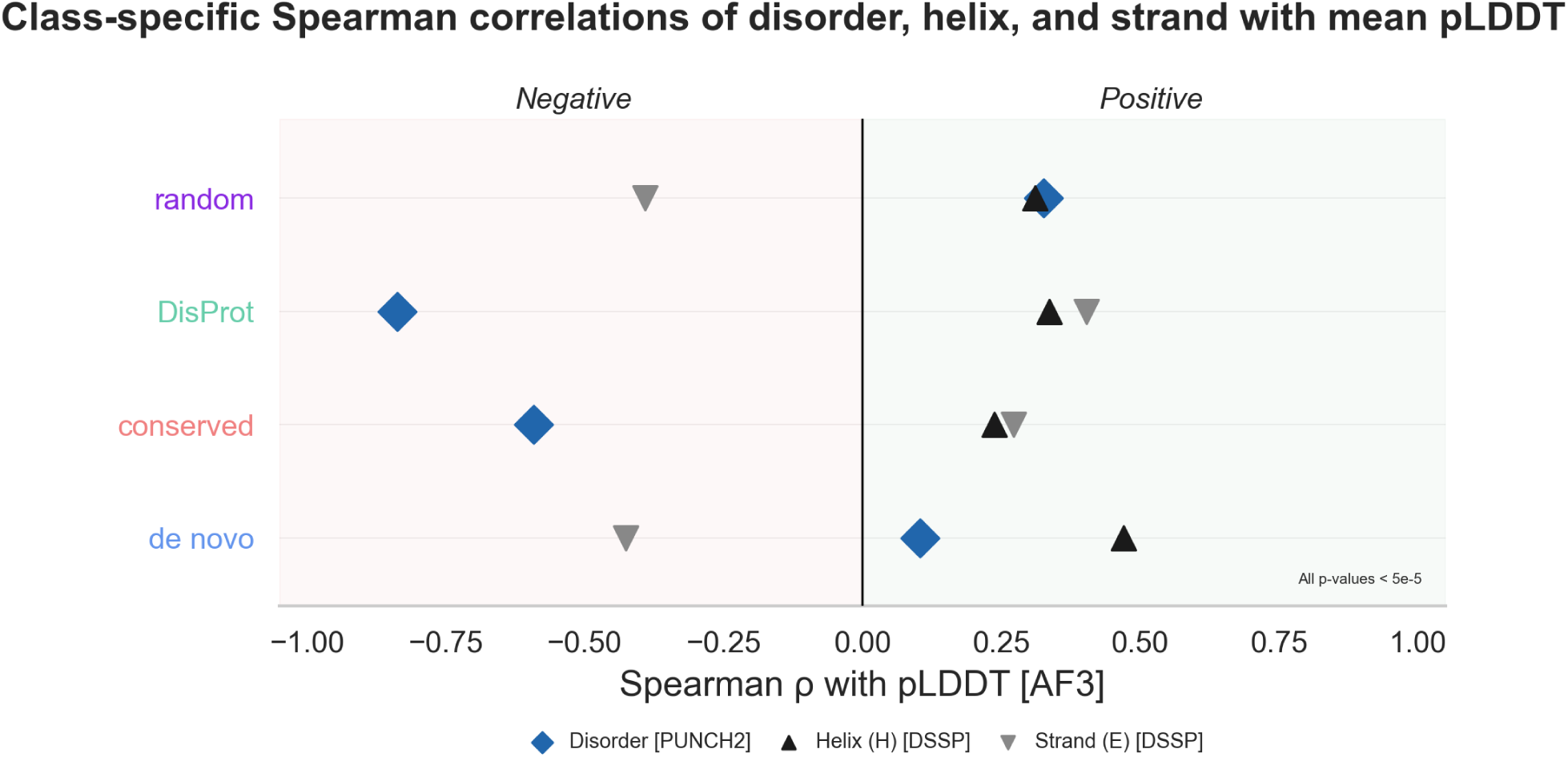
Class-specific correlations of disorder and secondary-structure features with AlphaFold3 pLDDT. Spearman correlations between mean per-protein AlphaFold3 pLDDT and PUNCH2 disorder, DSSP helix fraction and DSSP strand fraction are shown separately for random, DisProt, conserved and *de novo* proteins. Positive correlations are shown to the right of zero and negative correlations to the left. The pLDDT-disorder relationship is positive in random and *de novo* proteins, but negative in conserved and DisProt proteins. The pLDDT-strand relationship is negative in random and *de novo* proteins, but positive in conserved and DisProt proteins. All displayed correlations have *p* < 5 × 10^−5^.

We tested transmembrane (TM) helices as a possible mechanistic explanation, motivated by the high pLDDT typically assigned to membrane *α*-helices ^97,98^, the low PUNCH2 disorder of membrane segments and prevalence of TM helices in *de novo* emerged proteins ^33,50,66,67,99^. Excluding TM-annotated residues (predicted by Phobius ^74^) or all helical residues from the correlation calculation, and within-protein paired comparisons of TM vs. non-TM mean pLDDT (Supplementary Figures S19, S28, S30) modulated but did not abolish the inverted random and *de novo* correlation signals. TM content is therefore not the dominant explanation, and we explored more general structural-compositional drivers.

### 3.3 A class-specific score-defined driver subset captures the discordance

To localise the discordance to specific proteins rather than treating it as a class-wide property, we defined a per-protein driver score combining the three defining variables (mean pLDDT, mean disorder, strand fraction; see Methods: Equation 1). Proteins with driver score ≥ 1.0 within their class jointly displayed unusually high pLDDT, unusually high disorder, and unusually low strand content. This produced strikingly different driver fractions across classes: 24.5% of *de novo* (*n* = 614/2,506), 29.4% of random (*n* = 736/2,507), 5.1% of conserved (*n* = 114/2,224), and only 1.3% of DisProt (*n* = 27/2,066) proteins were classified as drivers, aligning with the observation that only *de novo* and random proteins show detectable discordance (Figure 3).

**Figure 3:**
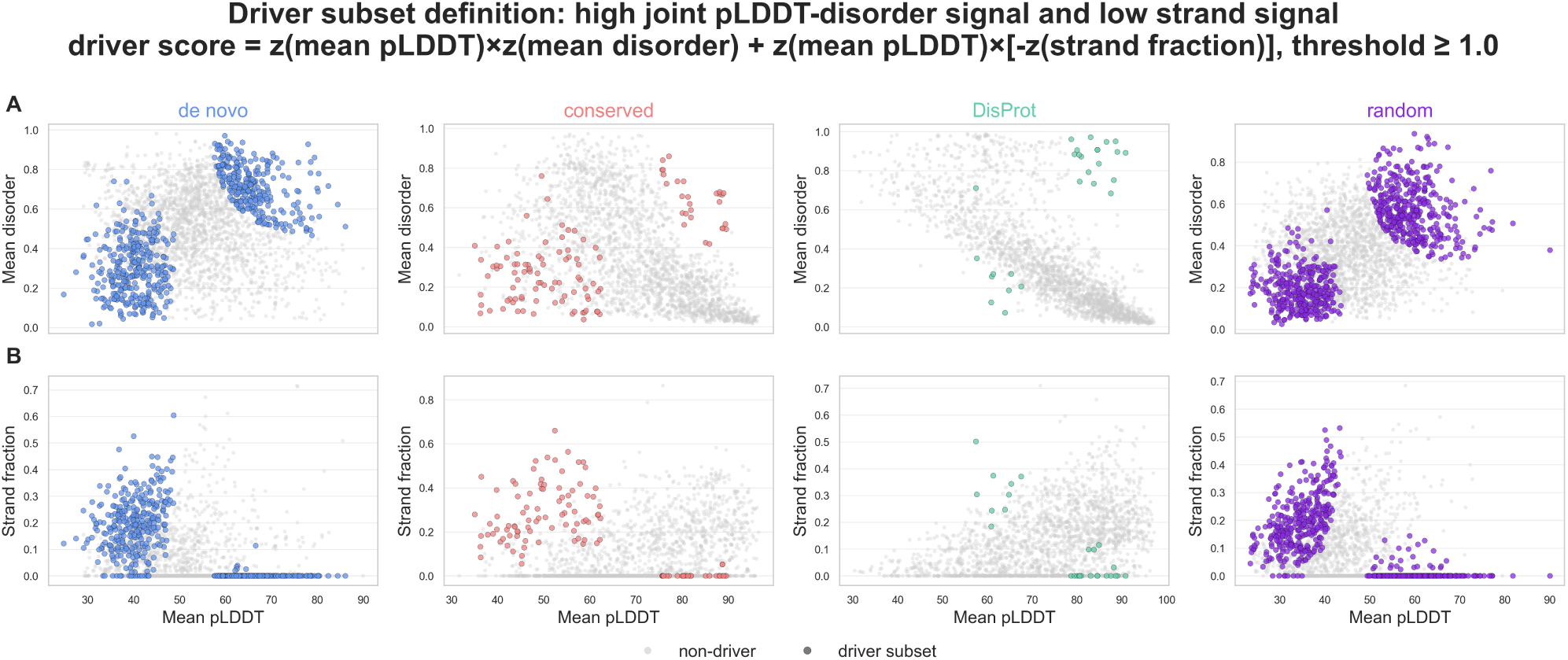
Definition of the score-defined driver subset. Proteins were classified as drivers using the class-standardised score *z*(mean pLDDT) × *z*(mean disorder) + *z*(mean pLDDT) × [−*z*(strand fraction)], with a threshold of ≥ 1.0. **(A)** Relationship between mean AlphaFold3 pLDDT and mean PUNCH2 disorder for each class. **(B)** Relationship between mean AlphaFold3 pLDDT and DSSP strand fraction for each class. Grey points indicate non-drivers and coloured points indicate driver-subset proteins. Drivers capture proteins with high joint pLDDT-disorder signal and low strand signal within each class.

Removing drivers normalised the class-level correlation pattern (Figure 4; Figure 5; Supplementary Figures S22-S32). For random sequences, pLDDT-disorder correlation collapsed from *ρ* = +0.33 (all proteins) to *ρ* = −0.09 after driver removal, while in driver-only proteins it rose to *ρ* = +0.68. For *de novo* proteins, pLDDT-disorder dropped from *ρ* = +0.12 to *ρ* = −0.26 after driver removal and reached *ρ* = +0.68 in drivers only. The strand-pLDDT correlation showed the same pattern: in random it weakened from *ρ* = −0.39 to *ρ* = −0.10 outside drivers (driver-only *ρ* = −0.71), and in *de novo* from *ρ* = −0.43 to *ρ* = −0.19 (driver-only *ρ* = −0.71). Conserved and DisProt classes, in which the canonical relationship already dominates, retained or strengthened their negative pLDDT-disorder correlation after driver removal (conserved *ρ* = −0.60 → −0.68; DisProt *ρ* = −0.85 → −0.87), and their small driver subsets reproduced the inverted signs found in the random/*de novo* drivers. The discordant pattern is therefore not a class-wide artefact but a localised phenomenon concentrated in a discrete subset of proteins.

**Figure 4:**
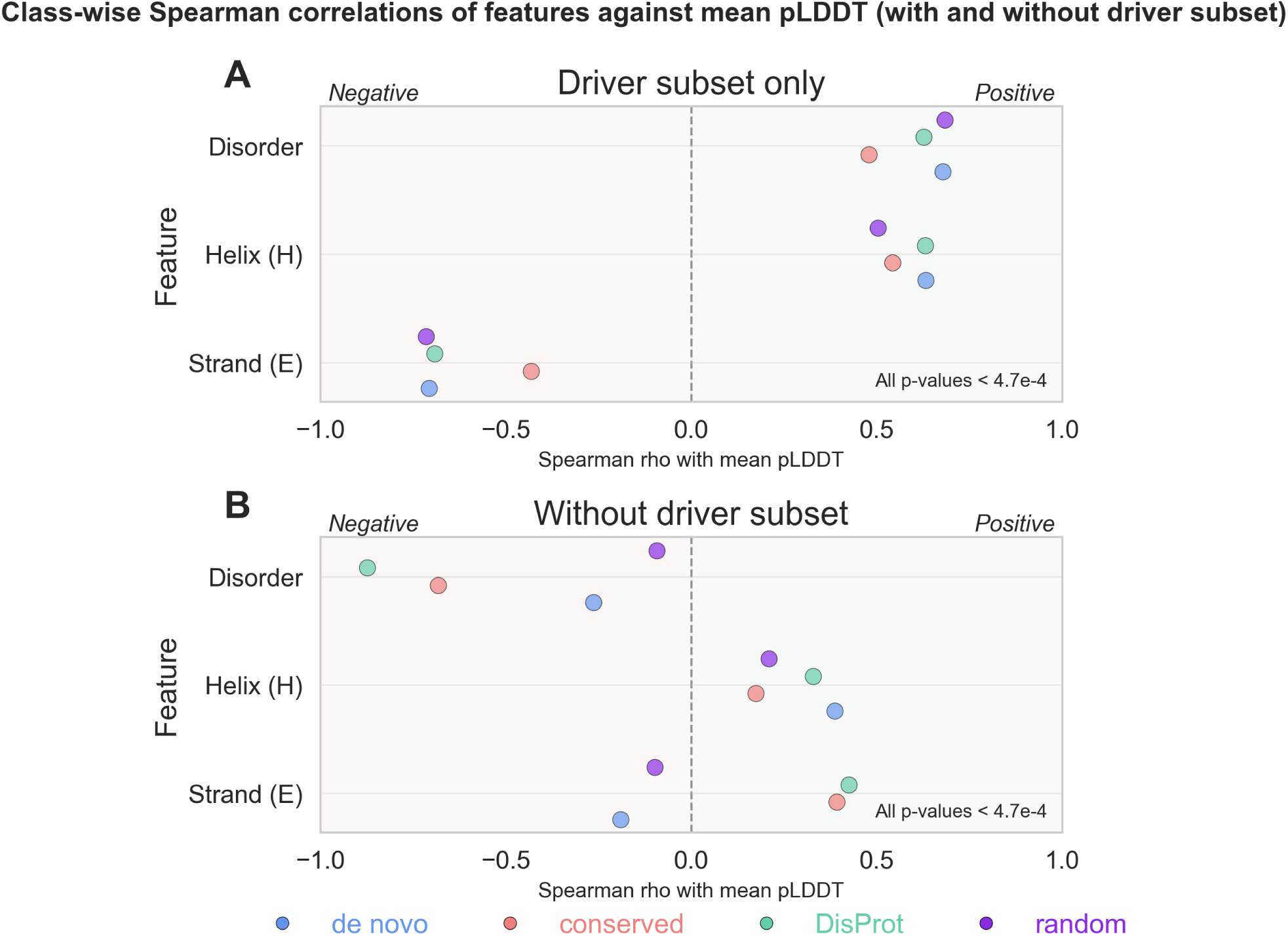
Driver-subset proteins concentrate the discordant pLDDT-feature correlation pattern. Class-wise Spearman correlations of mean AlphaFold3 pLDDT with PUNCH2 disorder, DSSP helix fraction and DSSP strand fraction were recalculated separately for driver-subset proteins and for proteins outside the driver subset. **(A)** Correlations within driver-subset proteins only. **(B)** Correlations after removing driver-subset proteins. In driver-subset proteins, pLDDT is positively correlated with disorder and negatively correlated with strand fraction. Removing drivers collapses or reverses the discordant correlations in random and *de novo* proteins and restores a more canonical relationship between pLDDT, disorder and secondary structure. All displayed correlations have *p* < 4.7 × 10^−4^.

**Figure 5:**
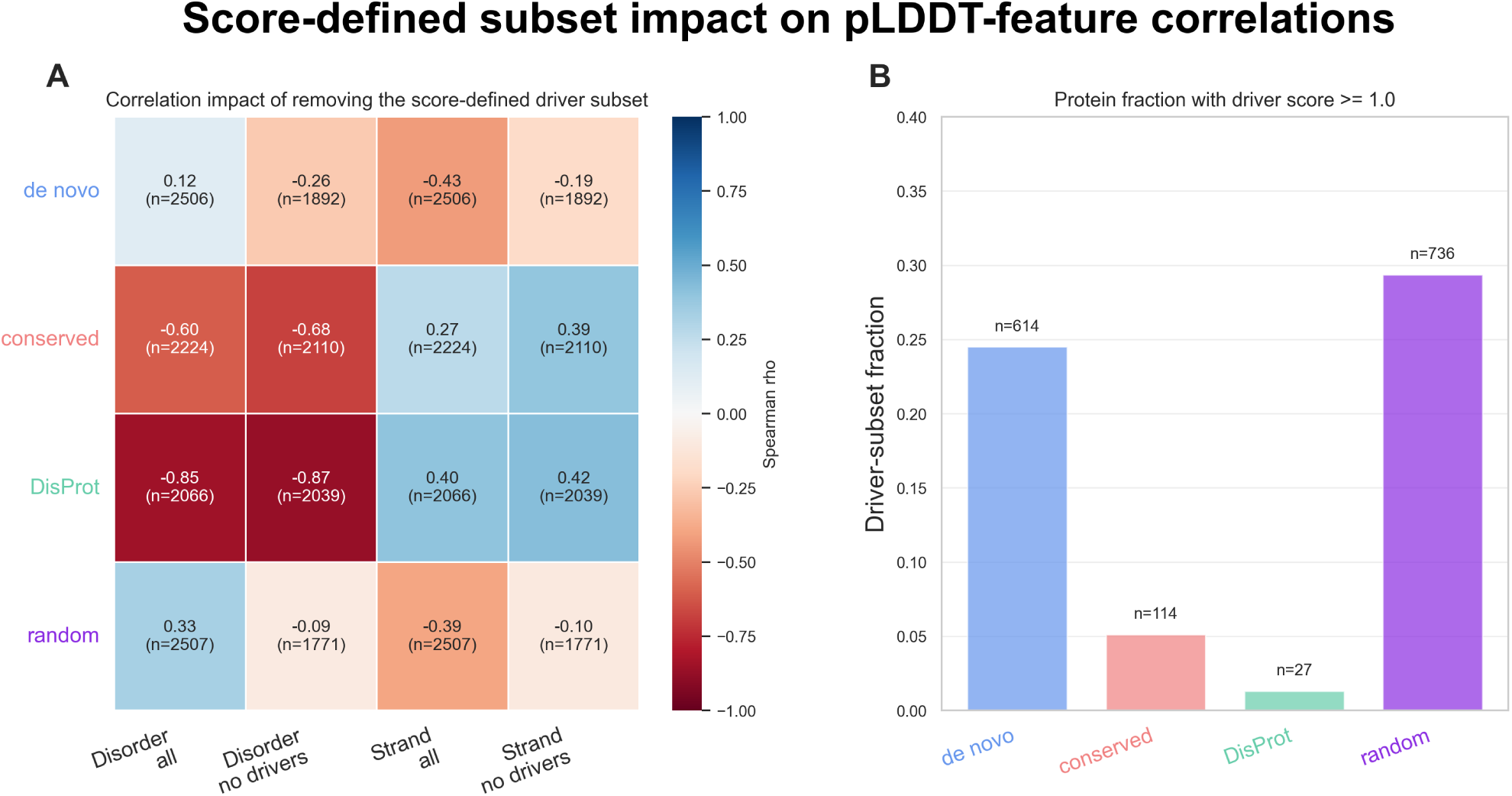
Impact of removing the score-defined driver subset on pLDDT-feature correlations. **(A)** Heat map of class-wise Spearman correlations between mean AlphaFold3 pLDDT and either PUNCH2 disorder or DSSP strand fraction, before and after driver-subset removal. Numbers inside cells give Spearman’s *ρ* and sample size. Removing the driver subset changes the pLDDT-disorder correlation from positive to negative in *de novo* and random proteins and weakens the negative pLDDT-strand relationship. **(B)** Fraction of proteins in each class classified as drivers using the score threshold ≥ 1.0. Driver-subset proteins are frequent in *de novo* and random proteins but rare in conserved and DisProt proteins.

### 3.4 Driver proteins differ from non-drivers in non-defining architectural and compositional features

Beyond the variables that defined them, driver proteins differ from non-drivers in a set of architectural and compositional properties that have direct biological interpretations (Figure 6; Supplementary Figures S33-S38). The single most consistent shared feature is enrichment in *β*-strand-bearing residues across all four classes (random: *δ* = +0.06, *q* = 8.5 × 10^−3^; *de novo*: *δ* = +0.24, *q* = 1.2 × 10^−24^; conserved: *δ* = +0.48, *q* = 2.0 × 10^−20^). Short, exposed *β*-strand stretches are well known to be structurally ambiguous: in isolation they often behave as transient or aggregation-prone elements rather than stable secondary structure ^100^, and in the absence of a co-folded *β*-sheet partner they sit on the boundary between order and disorder (a regime where AlphaFold-style models are calibrated against high-resolution PDB entries that under-represent transient strands and where disorder predictors - trained on residues annotated disordered in solution - flag the same residues as flexible). Drivers therefore appear to concentrate in proteins whose structural identity hinges on *β*-content that the two predictors interpret in opposite directions.

**Figure 6:**
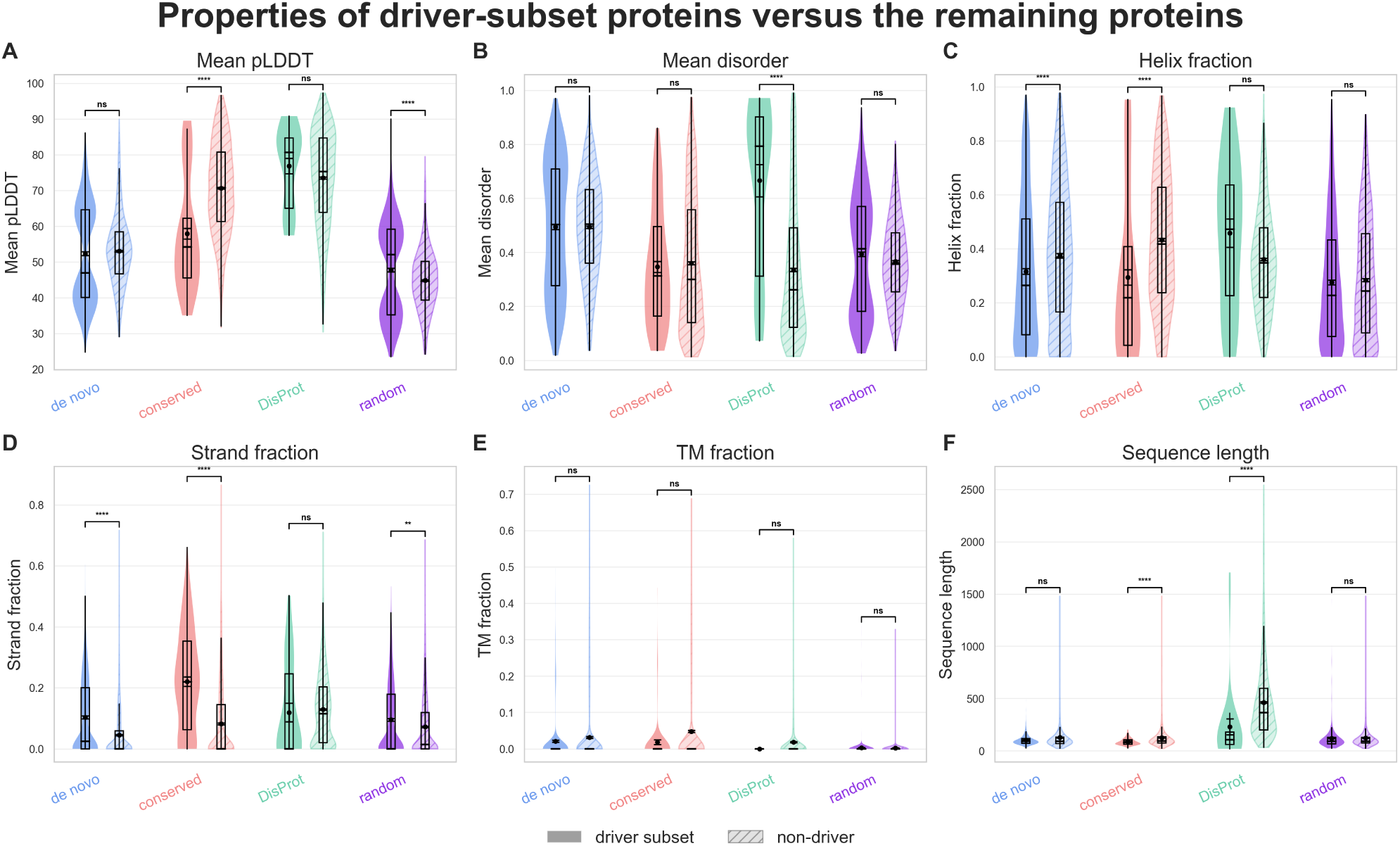
Properties of driver-subset proteins compared with non-driver proteins. Per-class distributions are shown for driver-subset proteins and non-drivers. **(A)** Mean AlphaFold3 pLDDT. **(B)** Mean PUNCH2 disorder. **(C)** DSSP helix fraction. **(D)** DSSP strand fraction. **(E)** Transmembrane-residue fraction. **(F)** Sequence length. Violin plots show the full per-protein distributions, with embedded box plots indicating median and interquartile range. Significance brackets compare drivers and non-drivers within each class. Driver proteins differ from non-drivers in class-specific architectural and compositional features, whereas transmembrane content does not consistently separate the two groups.

Drivers in the conserved and DisProt sets are additionally substantially shorter than their non-driver counterparts (conserved: median driver length 87 vs. non-driver 126, *q* = 1.8 × 10^−7^; DisProt: 230 vs. 463, *q* = 6 × 10^−8^) and depleted in *α*-helix content (conserved: *δ*_helix_ = −0.34, *q* = 3.5 × 10^−9^). Technically, structure and disorder predictions for very short proteins warrant particular caution, as short sequences provide limited structural and evolutionary context and frequently exhibit substantial conformational flexibility ^62–65^. Biologically, this is consistent with these drivers being short, helix-poor proteins or domains that are too small to bury a hydrophobic core, often characteristic of peripheral-or partner-binding modules whose structure depends on context. AlphaFold3 can still assign locally high pLDDT to such compact pieces, while PUNCH2 calls them disordered because in solution they are conformationally mobile until a binding partner is present. Crucially, drivers are not consistently enriched for transmembrane content (all |*δ*| ≤ 0.13), so this discordance cannot be explained by membrane *α*-helices, which would otherwise have been a parsimonious mechanistic candidate.

The class-specific compositional fingerprints are also biologically informative. *De novo* drivers are enriched in *β*-strand and depleted in helix relative to other *de novo* proteins, while *de novo* proteins generally contain more often helices (Figure 1;Figure 6). Random drivers occupy a similar architectural niche, indicating that the composition-length combination of *Drosophila de novo* sequences is sufficient to produce the same predictor disagreement without any selection. Conserved drivers, by contrast, are short, helix-poor, *β*-enriched proteins - exactly the architecture of compact accessory modules and small binding domains that adopt their fold conditionally. Ordinations on standardised non-defining feature sets show that the score-defined drivers occupy class-specific structured regions rather than a single universal cluster (Figure 7; Supplementary Figures S39, S43-S45), supporting the interpretation that several distinct biological architectures funnel into the same predictor-disagreement phenotype. We further examined the local sequence grammar of drivers using class-specific 3-mer and 5-mer enrichment within helical, strand-rich, and disorder-rich windows (Supplementary Figures S40-S44). Driver-enriched motifs are context dependent and class dependent, with distinct signatures in N-terminal vs. central windows and in helical vs. strand-rich vs. disorder-rich windows. The recurrent biological signal is enrichment in motifs combining hydrophobic and small/turn-prone residues at the local scale - the kind of compositional grammar that supports transient secondary structure or molten-globule character. The driver phenotype therefore has real local sequence content rather than reflecting only global protein-level averages, and its motif vocabulary is consistent with proteins whose *in vivo* structural state is conditional rather than fixed.

**Figure 7:**
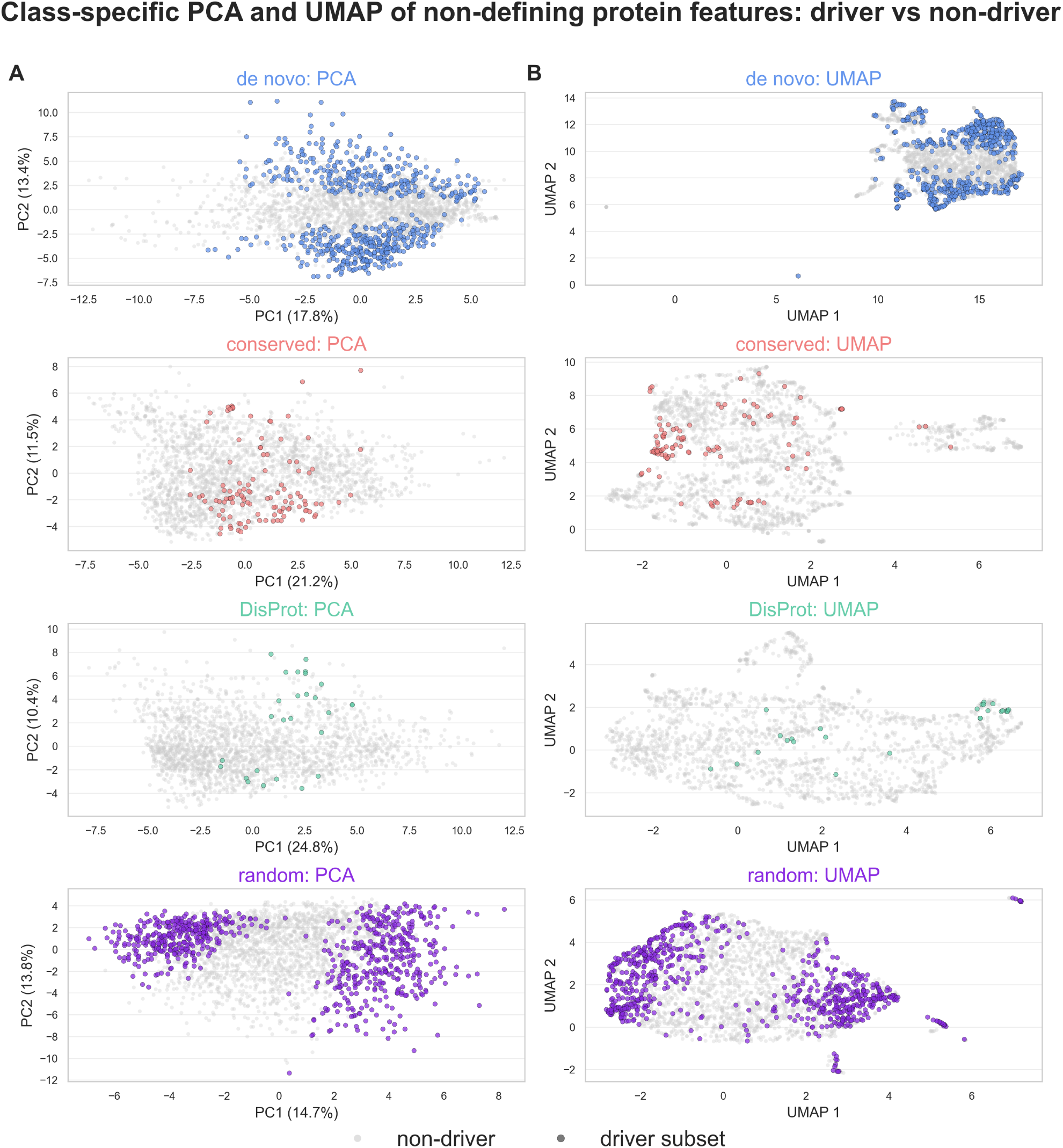
Ordination of driver and non-driver proteins using non-defining protein features. Principal component analysis and UMAP were performed separately for each class using protein-level features not directly used in the driver-score definition. **(A)** PCA ordinations for *de novo*, conserved, DisProt and random proteins. **(B)** UMAP ordinations for the same classes. Grey points indicate non-drivers and coloured points indicate driver-subset proteins. Driver proteins occupy structured, class-specific regions of non-defining feature space rather than forming a single universal cluster across all sequence classes.

### 3.5 A non-defining-feature classifier recovers driver proteins

To test whether drivers are also recoverable from features that do *not* include the defining variables, we trained class-specific histogram gradient-boosting classifiers using only architectural and compositional protein-level features (helix and coil fractions; TM and signal-peptide summaries; low-complexity and repeat fractions; Shannon entropy; hydropathy; charge; sequence length; regional helix/coil/TM/disorder fractions; per-amino-acid frequencies); pLDDT-, disorder-, and strand-derived variables were excluded (see Methods). Repeated stratified 3-fold cross-validation (20 repeats) gave class-wise mean average precision 0.62 in random and 0.58 in *de novo*, with ROC AUC ≈ 0.77-0.78; performance dropped on the lower-prevalence conserved (AP = 0.34, AUC = 0.82) and DisProt (AP = 0.29, AUC = 0.83) classes, consistent with strong label imbalance (1.3-5.1% drivers; Supplementary Figures S46-S50). Permutation importance identified helix and coil fractions, sequence length, global Shannon entropy, and hydropathy as the strongest non-defining predictors of driver status, with class-specific differences in which feature dominated (Figure 8; Supplementary Figures S46-S50). Score-defined and classifier-recognised driver subsets overlap but not completely, consistent with the interpretation that drivers form a compositionally enriched, class-dependent group rather than being fully separable (Supplementary Figure S49).

**Figure 8:**
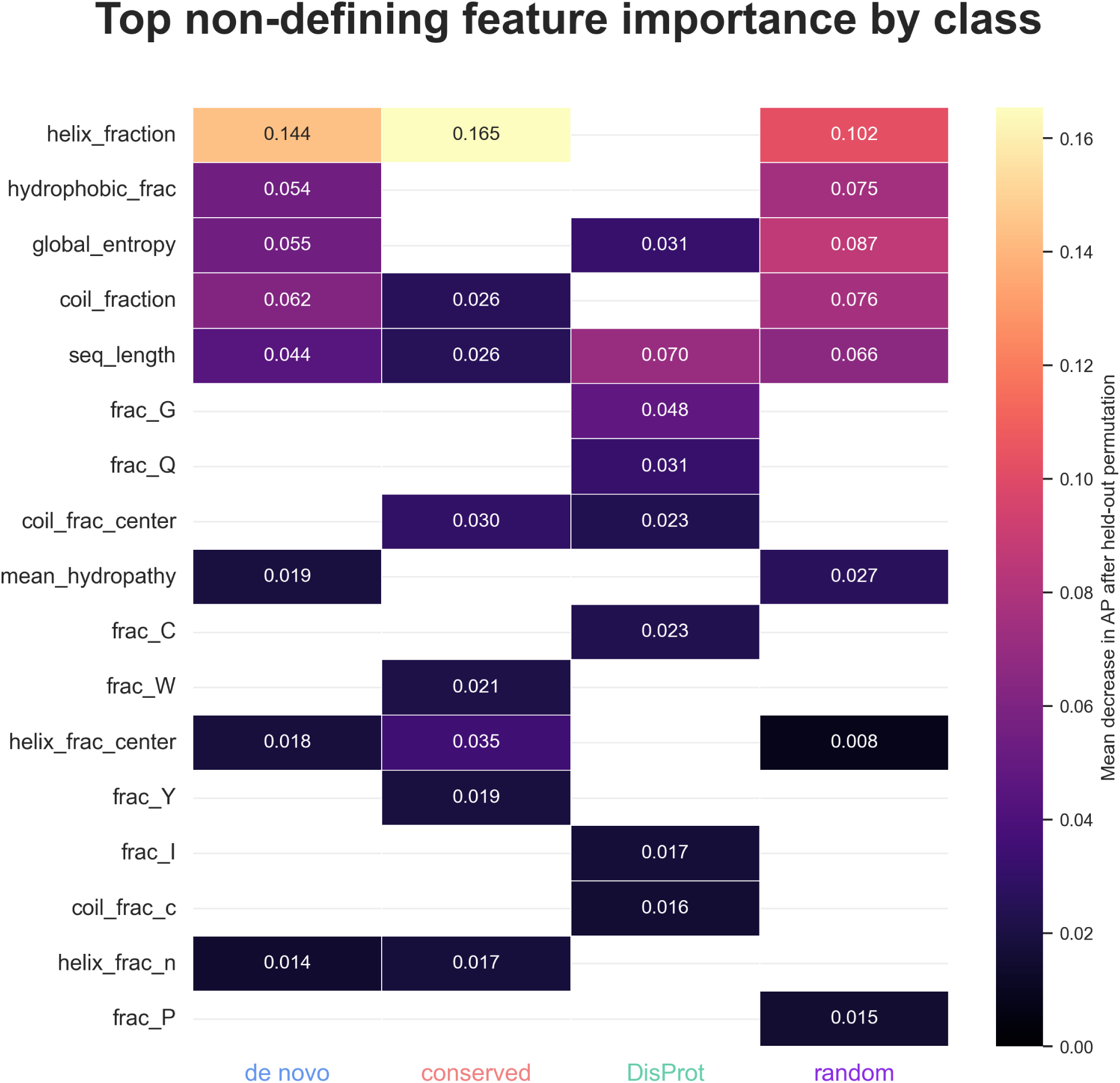
Non-defining features predictive of driver-subset status. Heat map of held-out permutation importance for the class-specific driver classifiers. Values indicate the mean decrease in average precision after permuting each feature in held-out data. Only features contributing to at least one class are shown. Helix fraction, hydrophobic residue fraction, global sequence entropy, coil fraction and sequence length are among the strongest non-defining predictors of driver status, with class-specific differences in feature importance across *de novo*, conserved, DisProt and random proteins.

## 4 Discussion

We set out to revisit a previously reported discordance between AlphaFold-style structure-prediction confidence and dedicated disorder predictions on *de novo* and random sequences ^32^. With the current best-of-class predictors AlphaFold3^21^ and the CAID-leading PUNCH2^22,25^ on identical sequence sets, the discordance not only persists but takes essentially the same form: pLDDT correlates positively with predicted disorder and negatively with *β*-strand fraction in random and *de novo* proteins, and oppositely in conserved and DisProt controls. The continued discordance with two independent and substantially upgraded predictor families argues against a model-specific calibration artefact and instead points to a property of the sequence space ^69^.

Our score-based localisation isolates this property to a discrete subset of proteins concentrated in the random (29.4%) and *de novo* (24.5%) classes but rare in conserved (5.1%) and especially DisProt (1.3%) proteins. Removing this subset normalises the class-level correlation structure, but the residual signals show that the effect is not perfectly discrete: positional context, sequence complexity, and compositional bias also modulate the predictor responses (Supplementary Figures S18-S21, S29). Our targeted test of transmembrane helices - a plausible mechanistic candidate, since TM helices typically receive high pLDDT and low PUNCH2 disorder - did not collapse the inverted correlations after exclusion of TM residues, indicating that membrane content is not the dominant explanation. The driver phenotype is more general, with helix and coil fraction, sequence length, entropy, and hydropathy emerging as the strongest non-defining predictors in our held-out classifier.

These observations have direct implications for protein engineering and *de novo* design. Modern design pipelines depend on AlphaFold-style scoring to verify that a designed sequence encodes the intended backbone ^36–38^, and disorder predictors are increasingly used to characterise naturally evolved *de novo* proteins, and viral orphans ^28,101^. Random and naturally emerged *de novo* sequences are useful stress tests for these tools because they sit far from the well-charted evolutionary regions on which the predictors were trained. Our results identify a concrete failure mode: a fraction of these sequences receives high AF3 confidence *and* high PUNCH2 disorder *and* low strand content together, a combination that is rare in evolved proteins and rare in DisProt. We do not interpret this as a generic claim that the predictors are wrong on these proteins - the underlying biophysical reality (folded? disordered? conditionally folded?) cannot be resolved from prediction alone - but as a quantitative warning that, on this specific subset, predictor outputs require explicit interpretation rather than direct trust. To this end, we provide a test suit for researchers to readily test their own AF3 predictions, in combination with disorder predictions, if their predictions are also discordant (see Methods).

Several limitations of our study should be noted. First, ground-truth structural data on naturally emerged *de novo* proteins remains very limited and exhaustive experimental validation is currently impractical at the dataset scale used here ^27^. Second, the random control set was constructed to match the *de novo* length and amino-acid distribution; alternative random regimes (e.g. uniform random, GC-content matched) might yield quantitatively different driver fractions, although the qualitative pattern of inverted correlations is unlikely to change given the consistency between random and *de novo* drivers in our analysis. Third, our analysis is restricted to monomeric structure prediction; AlphaFold3-Multimer and complex-aware analyses might modulate the high-pLDDT/high-disorder combination in some drivers. Finally, we did not benchmark protein language model based structure predictors such as ESMFold here, because we and others previously showed that ESMFold reproduces the same inverted pLDDT-disorder pattern on these sequence classes ^32,48^, and protein language model structure predictors did not match AlphaFold-style accuracy at CASP15 despite their reduced MSA dependence ^59^; adding such a predictor would not change the qualitative conclusion of this study, which is that the upgrade from AF2/flDPnn to AF3/PUNCH2 does not resolve the discordance.

While modern machine-learning models can generalize beyond their training data ^102–104^, prediction accuracy generally decreases as inputs become increasingly out-of-distribution relative to the data on which the models were trained. Orphan proteins occupy sparsely sampled regions of sequence space and therefore constitute precisely such out-of-distribution inputs for current protein structure prediction models ^52^. Extending reliable structure prediction into these unexplored regions of sequence space will therefore require experimentally determined structures from the “dark” regions of sequence space to expand the diversity of future training datasets ^52^. Expanding experimental coverage of the dark regions of sequence space will provide essential training data, but achieving true extrapolation beyond known proteins will likely also require models that increasingly capture the physical principles underlying protein folding rather than relying predominantly on statistical regularities in existing sequence–structure relationships ^105^.

In summary, the discordance between structure-prediction confidence and disorder prediction on *de novo* and random sequences is not resolved by AlphaFold3 and PUNCH2, persists with similar magnitude and sign as for AlphaFold2 and flDPnn, and is concentrated in a compositionally identifiable subset of proteins rather than being a uniform property of these classes. As structure and disorder predictors are pushed further into protein engineering and into the analysis of newly emerged proteins, this discordant subset is the part of sequence space where their outputs are most likely to mislead and where complementary biophysical or experimental evidence remains essential.

## Data and software availability

All sequence datasets (FASTA), AlphaFold3 rank-0 predictions (PDB), PUNCH2 disorder predictions (raw and per-protein/per-residue tables), DSSP annotations, Phobius predictions, complexity and repeat features, derived per-protein and per-residue master tables, statistical summary tables, score-defined driver subsets, classifier outputs, SI figures, and SI tables are deposited on Zenodo (DOI:10.5281/zenodo.22083740). Analysis scripts are available on GitHub (https://github.com/ArsLeicholt/protein-driver-discordance-AF3-PUNCH2). The standalone driver-assessment utility and usage documentation is deposited in a separate GitHub repository: https://github.com/ArsLeicholt/driver_assess.

## Acknowledgments

We thank Erich Bornberg-Bauer for access to computational resources. Predictions were performed on PALMA II HPC (subsidized by the Deutsche Forschungsgemeinschaft (INST 211/667-1)). This publication and other research outcomes are supported by the predoctoral program AGAUR-FI ajuts (2025 FI-3 00065) Joan Oró, of the Department of Research and Universities of the Generalitat of Catalonia, as well as the European Social Plus Fund.

## Author contributions

L.A.E. and L. M. conceived the study. L.A.E. performed the analyses, and drafted the manuscript. Both authors edited and approved the final manuscript.

